# Formulation of substrates with agricultural and forestry wastes for Camellia oleifera Abel seedling cultivation

**DOI:** 10.1101/2022.03.14.484217

**Authors:** Nianjin Wang, Fei Zhou, Jinping Zhang, Xiaohua Yao, Xiaofeng Zhang, Lingyan Zhan, Jieman Li

## Abstract

Five compound substrates for the cultivation of Camellia oleifera Abel seedlings were obtained by substituting the peats in conventional substrate formulas with the composts of Camellia oleifera shell, pine chips, palm fiber residues, chicken manure, and sheep manure. The changes in the physical and chemical properties of the substrate after seedling cultivation were determined and the effects of these substrates on the development of Camellia oleifera seedling were analyzed. It was found that the survival rates of the one-year-old seedlings produced from stem cuttings in all groups were greater than 97.5% after 6 months cultivation. The seedlings exhibited greater seedling heights, larger ground diameters, longer root systems and larger total root volume than those obtained with conventional substrates. The compound substrate formulated with the compost of Camellia oleifera shell+ palm fiber residue+ chicken manure (A3), vermiculite and perlite (6:3:1) is the most optimal, which gives the 100% survival rate of cultivated seedlings, the greatest seedling heights, and the largest ground diameters. In particular, the ground diameters and 26.67% of the seedling heights reach the grade 1 standard for the two-year-old seedlings.

## 1. Introduction

In China, 2 billion tons of a variety of agricultural and forestry wastes are produced every year. These wastes are renewable and biodegradable biomasses, yet only a small portion are reused. The majorities of them are incinerated, buried, or disposed randomly, which is the waste of resources and causes environmental pollutions (Guo et al., 2021; Zhao et al., 2021; Lu et al., 2021). Agricultural and forestry wastes can be reutilized as the cultivation substrates for flowers, seedlings, and vegetables after treated by high-temperature aerobic fermentation (Zhao et al., 2021). Such applications not only re-utilize biomass wastes and reduce environmental pollutions, but also effectively improve the ecological environment (Nie, 2017) and realize carbon sequestration.

Camellia oleifera Abel. is an important woody oil tree species in southern China (Zhuang, 2008). The rapid development of tea oil processing industry has been dramatically increasing the production of by-products, such as Camellia oleifera shells. Composting of these by-products can not only prevent the waste of resources and environmental pollution, but also provide plant cultivation substrates and organic fertilizers to improve soil quality and inhibit the occurrence of soil-borne diseases. Seed box method with light substrates shows great advantages, such as all-season operation, high survival rates, no slow growth period, fast growth, less root damage during transplanting, long afforestation seasons, and low transportation costs for seedling production. (Wang et al., 2018; Liu et al., 2019; Lv et al., 2017; Yang et al., 2018). Substrate selection is one of the key factors affecting the performance of seed box production of seedling (She et al., 2020). The conventional camellia oleifera seedling production is mostly based on the substrates formulated with topsoil and peat. Peat is a non-renewable resource, and its efficiency for Camellia oleifera seedling production can be further improved (Wu et al., 2017). Therefore, substituting peat with agricultural and forestry wastes has become a research hotspot. For example, Lu et al. reported the Camellia oleifera seedling production on the substrate formulated with the compost of forest wastes, weeds, and vermicompost (Lu et al., 2021). Wu et al. successfully obtained Camellia oleifera seedlings with the substrate containing the compost of sawdust, decomposed litter, biogas residue and yellow soil (Wu et al., 2017). Dai et al. also formulated the substrate with the compost of bagasse, cassava skin, peanut shells, charcoal ash and garden soil for the production of Camellia oleifera seedlings (Dai et al., 2016). All these reports used the composts of agricultural and forestry wastes as the substrates for the seedling production of Camellia oleifera, yet lack the detailed information of the sources of the raw materials, composting formula and composting method. In addition, the variation in the sources of substrate raw materials makes it difficult to obtain the consistent composts. The quality of the compost of Camellia oleifera shell has been successfully improved by adding different nitrogen sources, such as urea, compound fertilizer, and pig manure and EM bacteria (Luo, 2011), but no application research on the composts has been carried out.

Composting is a major processing process of agricultural and forestry wastes. It is generally divided into aerobic composting and anaerobic composting. Aerobic composting decomposes organic matters under aerobic conditions mostly into CO_2_ and water and releases heat. Anaerobic composting mainly produces methane, CO_2_ and low molecular weight intermediates, such as organic acids, etc. Most conventional composting is conducted under anaerobic conditions, which tend to release odors (Li et al., 2017). The modern composting process is mainly conducted under aerobic conditions with controlled water contents, C/N ratios and ventilation. Aerobic composting can transform unstable agricultural and forestry waste into stable substances, during which pathogenic bacteria are killed. The composting thus can be safely handled and stored. They are also a good seedling substrate, soil conditioner and organic fertilizer (Zhang et al., 2021).

In this work, aerobic composting was conducted using camellia oleifera shells, pine chips, palm fiber residues, chicken manure and sheep manure as the major raw materials. The composts were respectively mixed with boiler slag, vermiculite, and perlite at certain ratios as the substrates for the cultivation of Camellia oleifera seedlings. The changes in physical and chemical properties of the substrates after seedling cultivation and the relationship between composting raw materials and the seedling development were comprehensively evaluated and analyzed, aimed to optimize a composting formula and substrate formula for the cultivation of Camellia oleifera seedlings and provide the research data and scientific reference for the recycling and composting of agricultural and forestry wastes, such as camellia oleifera shells, as the substrate exclusively for the seed box cultivation of Camellia oleifera seedlings.

## 2. Materials and methods

### 2.1 Compost materials and preparations

Camellia oleifera shells, palm fiber residues, chicken manure, pine chips, and sheep manure were provided by Anhui Wanxiuyuan Ecological Agriculture Group Co., Ltd., (Anqing, Anhui Province, China). The Camellia oleifera shells and pine chips were smashed to the sizes of 5-8 mm before use. The compost bacteria were purchased from Yijiayi Biological Engineering Co., Ltd. (Zhengzhou, China) and urea was obtained from Henan Jinkai Chemical Investment Holding Group Co., Ltd (Zhengzhou, China). The primary properties of the compost raw materials are listed in Table 1.

**Table 1.** Primary properties of compost raw materials

| Raw material/Property | TOC (%) | TN (%) | C/N | Cellulose/% | Hemicellulose/% | Lignin/% |
| --- | --- | --- | --- | --- | --- | --- |
| *Camellia oleifera* shells | 51.80±2.58 | 0.729±0.03 | 71.359±3.74 | 18.62±0.22 | 49.34±0.07 | 29.71±0.14 |
| Brown silk chips | 46.80±2.23 | 2.68±0.14 | 17.557±1.76 | 28.15±0.18 | 20.58±0.08 | 44.08±0.16 |
| Pine chips | 54.20±2.74 | 0.285±0.01 | 190.964±5.19 | 48.36±0.11 | 27.59±0.12 | 19.58±0.21 |
| Chicken mature | 34.39±1.75 | 2.73±0.14 | 12.667±1.31 | - | - | - |
| Sheep mature | 36.47±1.81 | 2.78±0.14 | 13.186±1.32 | - | - | - |
| Urea | 20±0.18 | 46.4±1.22 | 43.1±1.12 | - | - | - |

“-” indicates extremely low content or none

### 2.2 Composting process

Composting was conducted in the composting plant of Anhui Wanxiuyuan Ecological Agriculture Group Co., Ltd. from August to November 2020 by the windows composting system. Four composting groups including A1: Camellia oleifera shell + urea (C/N 60); A2: Camellia oleifera shell+ sheep manure (C/N 55); A3: Camellia oleifera shell+ palm fiber residue + chicken manure (C/N 55) /N 30); and A4: pine chips+ chicken manure (C/N 40.7) were mixed and stirred evenly. The moisture content of each pile was adjusted to 55-60% and 0.1% fermentation bacteria were added to each pile. Each pile was then thoroughly mixed, stirred evenly, and stacked for high-temperature aerobic fermentation at the same time. The composting piles were turned for aeration and sampled by the five-point sampling method every 7 days. For each sample, half portion was stored at −20°C and the other half was dried at 65°C and ground.

### 2.3 Compound substrate preparation and seedling nursing

Seedling nursing was carried out at the nursery of Anhui Wanxiuyuan Ecological Agriculture Group Co., Ltd. located in Taihu County, Anqing, Anhui Province between 30°09’ N to 30°46’ N and 115°45 ‘E to 116°30 ‘E. The temperatures during the nursing varied between from −6 °C and 34 °C, and the temperature lower than 0 °C lasted for 24 days.

The compound substrates were prepared with the composts obtained above and inorganic matrices using the formulas shown in Table 2. In December 2020, one year old Camellia oleifera seedlings with intact root systems, the heights of 13 ± 3 cm and ground diameters of 0.22 ± 0.02 cm were respectively transplanted into seed boxes with the diameter of 16 cm and height of 16 cm. The boxes were filled with the compound substrates listed in Table 2. For each substrate, 200 replicates were prepared and allowed to grow for 6 months. The substrates were then sampled for analysis. During the cultivation, the substrates were maintained moist, and watered thoroughly when the substrate became dry during the fast growing period.

**Table 2.**
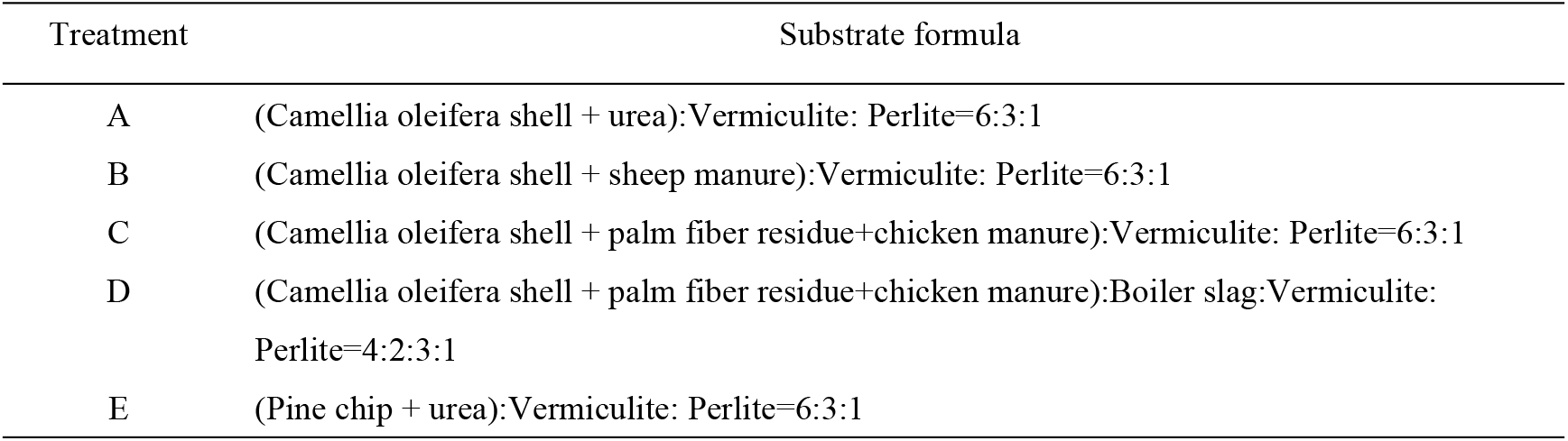
Formulas of compound substrates

### 2.4 Analysis methods

During composting, the compost temperature was measured in the upper (10 cm from the top), middle and lower (10 cm from the bottom) parts of the pile at around 3:00 pm every day, and the ambient temperature of the day was recorded at the same time. The bulk density, total porosity, aeration porosity, water-holding porosity, electrical conductivity (EC) and pH were also measured following the Chinese Forestry Industry Standard GB/T 33891-2017 of organic substrates for greening. Total organic carbon (TOC), total nitrogen (TN), total phosphorus (TP), total potassium (TK) and germination index (GI) were determined by the method of Zhang et al. (2018). The organic matter content, total nutrient content, C/N ratio and survival rate of Camellia oleifera seedlings were calculated with Eq. (1), (2), (3) and (4), respectively.

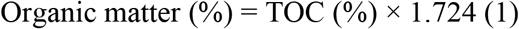

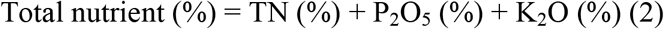

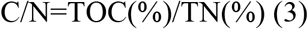

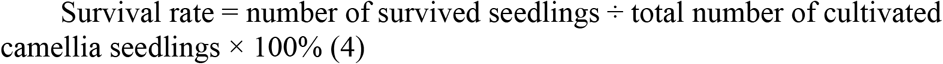

After 185 days of cultivation, 60 seedlings in each group were randomly collected and measured for ground diameters and heights. Five seedlings were randomly selected from each group and analyzed for the lengths, surface areas, volumes and average diameters of their root systems using a Wanshen LA-S root system analyzer to evaluate the effects of the physical and chemical properties of substrate on the growth of Camellia oleifera seedling. The obtained data were processed and analyzed using the EXCEL and SPSS softwares.

## 3. Results and discussion

### 3.1 Compost temperature

Temperature is one of the important indicators to evaluate the compost maturity (Gu et al., 2015; Bustamante et al., 2008). The composting process is generally divided into four phases, e.g., the mesophilic phase, the thermophilic phase (>50 °C), the cooling phase, and maturation phase. Pathogenic microorganisms, insect eggs and weed seeds can be killed at the thermophilic phase when the compost temperatures increases to over 55 °C, producing harmless composts. The fermentation bacteria are rapidly deactivated as temperature increased to over 63 T, leading to temperature drops of the compost (Bernal et al. 2009). As can be seen from Figure 1, the temperatures of all composting piles are much higher than the ambient temperature during the composting, indicating that the ambient temperature has little effects on the composting. The initial temperatures of the 4 groups of composting piles were 27±0.6°C. The temperatures A1, of A2, A3 and A4 composting piles increased to above 50°C on the day 6, 7, 3 and 3, respectively, and remained at 50°C for 20, 23, 39, and 9 days, and at 60 °C for 4, 7, 22, and 6 days, respectively, complying the safety and hygiene requirements (Wang et al., 2017). The four composting piles entered the cooling phase on day 33, 34, 44 and 15, respectively and their maturation phases lasted 38, 42, 44 and 21 days, respectively. The sudden temperature drops are due to heat dissipation during manual turning for aeration (Qian et al, 2014; Guardia et al, 2010a).

**Figure 1.**
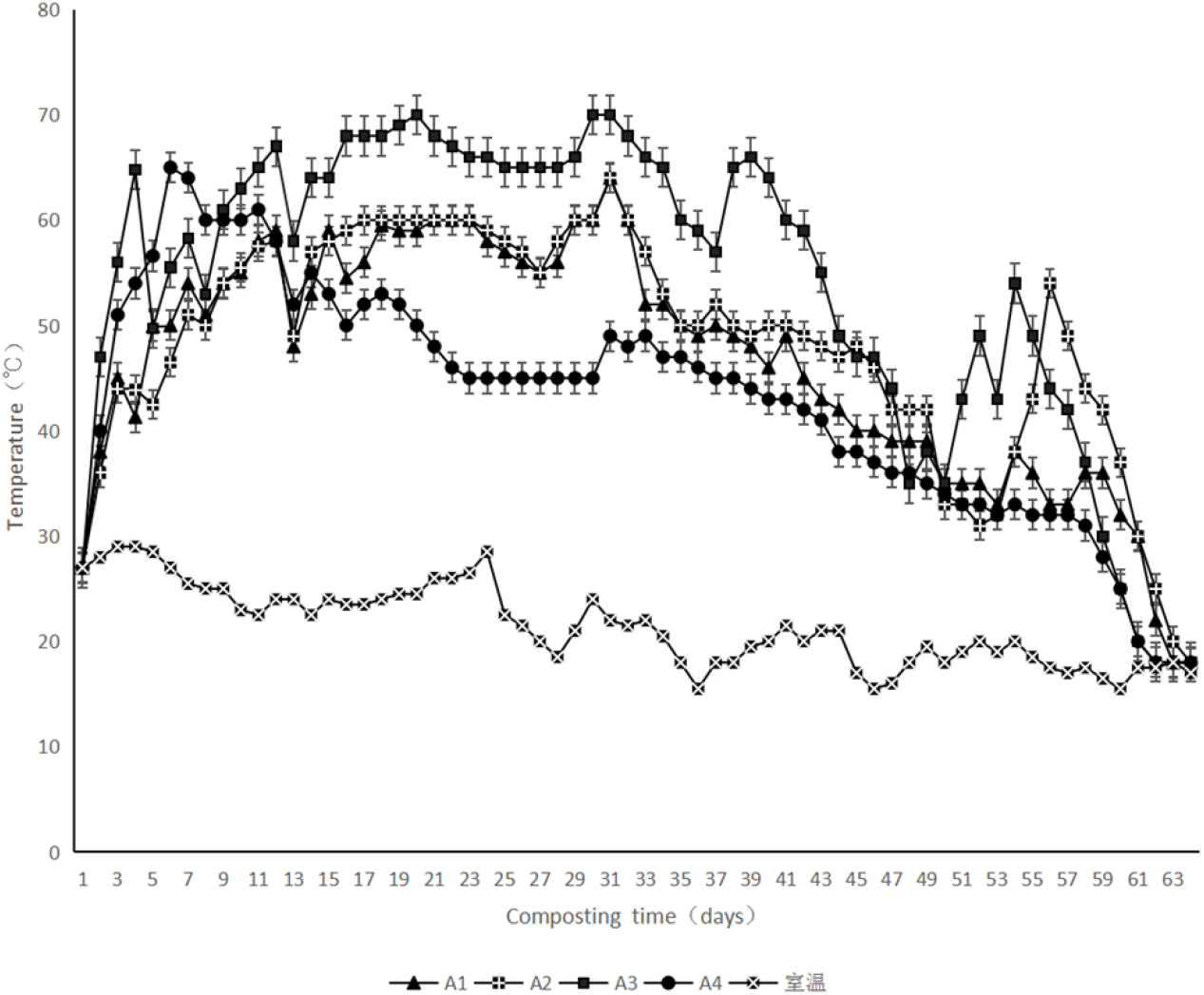
Temperature changes of the four composting piles during composting

The A3 composting pile exhibited the largest heating rate, the highest high temperature, and the longest thermophilic phase. It can be explained that, first, the initial C/N (30) of A3 is most suitable for microbial reproduction (Bernal et al. 2009). Second, there is a large amount of chicken manure in A3. Chicken manure is rich in protein and urea that can be easily decomposed and utilized by microorganisms. Third, the palm fiber residues can improve the aeration porosity of the compost, which is conducive to the proliferation of the microorganisms and heat generation. Groups A1, A2 and A3 all contain camellia oleifera shells, but have different C/N ratios due to different excipients. The C/N ratio of A3 is 30, which is in the optimal range of 25-35, while those of A1 and A2 are 60 and 55, respectively, suggesting insufficient nitrogen. The activity of microorganisms is low under the insufficient nitrogen conditions, which slows down the composting reaction (Ma et al., 2020). The temperature of A4 increased rapidly and thus its initial temperature was higher. However, it entered the cooling phase earliest and the temperature dropped fastest. The nutrients in the chicken manure are first decomposed rapidly. The hemicellulose content in the pine chip is only 27.59%, and the majority is cellulose and lignin that are difficult to be degraded. In contrast, the Camellia oleifera shells in other three groups are rich in easily degradable hemicellulose (49.34%), which contributes to the long thermophilic phases. These results suggest that the physical and chemical properties of raw materials directly decide the composting process and the physical and chemical properties of compost (Table 3).

**Table 3.** Physical and chemical properties of the composts of agricultural and forest waste and other components for the preparation of compound substrates

| Group | Moisture<br>content MC/% | bulk density $\text{g}\cdot\text{cm}^{-3}$ | Total porosity % | Aeration<br>porosity % | Water-holding<br>porosity % | pH | EC ($\text{ms}\cdot\text{cm}^{-1}$) |
| --- | --- | --- | --- | --- | --- | --- | --- |
| A1 | 10.27±0.52 | 0.39±0.018 | 73.94±3.68 | 6.07±0.29 | 67.87±3.39 | 7.00±0.21 | 0.13±0.02 |
| A2 | 9.46±0.48 | 0.40±0.021 | 74.13±3.70 | 9.30±0.47 | 64.83±3.23 | 8.00±0.26 | 0.08±0.03 |
| A3 | 13.82±0.68 | 0.47±0.024 | 65.90±3.31 | 11.83±0.60 | 54.07±2.71 | 7.79±0.24 | 0.20±0.01 |
| A4 | 22.47±1.11 | 0.42±0.02 | 83.40±4.16 | 1.80±0.089 | 81.60±4.07 | 7.00±0.22 | 0.06±0.03 |
| Group | TN% | TK% | TP% | TOC% | total nutrient % | GI% | organic matter % |
| A1 | 1.07±0.05 | 1.83±0.09 | 0.37±0.02 | 38.20±1.92 | 3.27±0.15 | 89.12±4.44 | 65.86±3.25 |
| A2 | 1.16±0.057 | 2.07±0.12 | 0.62±0.03 | 36.30±1.82 | 3.85±0.21 | 90.13±4.49 | 62.58±3.11 |
| A3 | 2.75±0.14 | 2.55±0.11 | 1.02±0.05 | 29.80±1.48 | 6.32±0.33 | 99.42±4.98 | 51.38±2.54 |
| A4 | 0.84±0.04 | 0.54±0.03 | 0.42±0.02 | 47.30±2.37 | 1.80±0.08 | 91.31±4.52 | 81.55±4.06 |
| Raw materials | Moisture<br>content MC/% | bulk density $\text{g}\cdot\text{cm}^{-3}$ | Total porosity % | Aeration<br>porosity % | Water-holding<br>porosity % | pH | EC($\text{ms}\cdot\text{cm}^{-1}$) |
| Boiler slag | 2.08±0.11 | 0.70±0.03 | 63.10±3.16 | 0.40±0.019 | 62.7±3.14 | 8.00±0.025 | 0.22±0.012 |
| Vermiculite | 1.55±0.078 | 0.13±0.01 | 95.04±4.75 | 30.03±1.51 | 65.01±3.24 | 7.00±0.021 | 0.047±0.0024 |
| perlite | 0.14±0.007 | 0.16±0.01 | 60.33±3.03 | 29.51±1.48 | 30.82±1.55 | 7.00±0.019 | 0.022±0.0011 |

### 3.2 Physical and chemical properties of composts and substrates

The composts of agricultural and forestry wastes can be used alone as a seedling substrate or can be formulated with other matrices as seedling substrates. The substrates with large bulk densities are favorable for the seedling handling. The cohesions of the substrates with small bulk densities are poor, which is inconducive to the fixation of root system (Li et al., 2020). The bulk densities of A1, A2, A3, and A4 composts are 0.39 g·cm^-3^, 0.40 g·cm^-3^, 0.47g·cm^-3^, and 0.42 g·cm^-3^, all within the ideal bulk density range of 0.1-0.8 g·cm^-3^ (Wang et al., 2021). Porosity is another important physical property that affects the aeration, drainage and water holding functions of a substrate. The total porosities of A1, A2 and A3 composts are 73.94%, 74.13% and 65.90%, respectively, which are all within the ideal total porosity ramge of seedling substrate (40%-75%) (Wu, 2006; Wang et al., 2013). The total porosity of A4 compost is 83.4%, which needs to be adjusted with other matrices. All the pH, seed germination indexes (GI), and organic matter contents of the composts are 7-8, 89.12-99.42, and 65.86-81.55%, respectively, all of which meet the standard NY/T525-2021 for organic fertilizer defined by the Ministry of Agriculture and Rural Affairs of the People’s Republic of China (pH 5.5-8.5, GI≥70%, and organic matter content≥30%). GI is a parameter that can be used to quickly evaluate the phytotoxicity of compost in a short period of time and is of significant importance in actual productions. The GI of the 4 groups of composts are >85%, and thus the composts can be considered non-toxic to plants (Yamamoto, N. et al. 2011). The total nutrient contents of A1, A2, A3, and A4 composts are measured to be 3.27%, 3.85%, 6.32%, and 1.80% respectively. The total nutrient content of A3 reaches the standard NY/T525-2021 for organic fertilizer (≥4%) and can be used as an organic fertilizer or seedling substrate raw material. The rest 3 groups can be used as cultivation substrates or substrate raw materials.

### 3.3 Changes in physical and chemical properties of compound substrates after seedling cultivation

The composts of agricultural and forestry wastes have relatively poor water holding capacities, and thus vermiculite with a better water holding ability is introduced. The raw materials listed in Table 3 are mixed at the ratios listed in Table 2, and the physical and chemical properties of the compound substrates are shown in Table 4. The bulk densities of compound substrates A, B, C, D, and E are measured to be 0.41 g·cm^-3^, 0.47 g·cm^-3^, 0.41 g·cm^-3^, 0.42 g·cm^-3^, and 0.41 g·cm^-3^, respectively and their total porosities are 73.83%, 67.83%, 66.70%, 68.17% and 74.60%, respectively, all within the ranges of 0.1-0.8 g·cm^-3^ and of 40%-75% (Wang et al., 2021; Wu, 2006; Wang et al., 2013). Their aeration porosities (29.31%, 28.12%, 25.44%, 24.22% and 23.10%), pH (7.03-7.43) and EC values (0.12-0.20 ms·cm^-1^) all conform to the national standard of the People’s Republic of China of “Organic Substrates for Greening” (GB/T 33891-2017) with the requirements of total porosity≥20%, pH 5.0-7.6 and EC≤0.65 ms·cm^-1^. All the pH do not conform to the Forestry Industry Standard of the People’s Republic of China LYT 2314-2014 “Technical Regulations for Seed Box Production of Camellia Container Seedling” (pH 5.0-6.5).

**Table 4.** Changes in the physical and chemical properties of compound substrates after seedling cultivation

| group |  | A | B | C | D | E |
| --- | --- | --- | --- | --- | --- | --- |
| bulk | before | 0.41±0.02 | 0.47±0.02 | 0.41±0.02 | 0.42±0.02 | 0.41±0.02 |
| density | after | 0.57±0.027 | 0.76±0.037 | 0.59±0.028 | 0.59±0.029 | 0.60±0.031 |
| g/cm<sup>3</sup> | increase % | 39.02±4.39 | 61.70±2.77 | 43.90±2.17 | 40.48±2.02 | 46.34±2.20 |
| total | before | 73.83±3.65 | 67.83±3.34 | 66.70±3.37 | 68.17±3.37 | 74.60±3.68 |
| porosity % | after | 58.07±2.86 | 58.80±2.94 | 61.40±3.17 | 64.50±3.29 | 59.60±3.95 |
|  | decrease | 21.35±1.03 | 13.31±0.63 | 7.95±0.36 | 5.38±0.22 | 20.11±0.95 |
| aeration | before | 29.31±1.35 | 28.12±1.31 | 25.44±1.22 | 24.22±1.19 | 23.10±1.12 |
| porosity % | after | 52.11±2.33 | 54.21±2.34 | 51.91±2.19 | 44.22±2.16 | 42.18±2.02 |
|  | increase | 77.79±3.82 | 92.78±4.55 | 104.05±5.07 | 82.58±3.92 | 82.60±3.85 |
| Water- | before | 44.52±2.21 | 39.71±1.94 | 41.26±1.99 | 43.95±2.08 | 51.50±2.42 |
| holding | after | 5.96±0.22 | 4.60±0.21 | 9.49±0.42 | 8.30±0.37 | 17.41±0.83 |
| porosity % | decrease | 86.61±4.25 | 88.42±4.31 | 77.00±3.68 | 81.11±3.94 | 66.19±3.20 |
| pH | before | 7.43±0.29 | 7.38±0.24 | 7.03±0.26 | 7.34±0.27 | 7.20±0.23 |
|  | after | 7.32±0.34 | 7.45±0.36 | 7.65±0.37 | 7.48±0.36 | 7.61±0.37 |
|  | increase% | -1.48±0.067 | 0.95±0.041 | 8.82±0.41 | 1.91±0.094 | 5.69±0.26 |
| EC | before | 0.15±0.002 | 0.15±0.002 | 0.20±0.002 | 0.12±0.003 | 0.12±0.003 |
| (ms·cm<sup>-1</sup>) | after | 0.46±0.023 | 0.22±0.012 | 0.24±0.007 | 0.18±0.005 | 0.24±0.002 |
|  | increase% | 206.66±2.0 | 46.00±2.3 | 20.00±1.4 | 50.00±2.3 | 50.00±1.33 |

Table 4 and Table 5 compare the physical and chemical properties of the five compound substrates before and after the seedling cultivation. As can be seen, the bulk densities, aeration porosities and EC values of all groups increase, while their total porosities and water-holding porosities, organic matter contents and total nutrient contents decrease after 6 months cultivation. It can be explained that, first, most of the hemicellulose in the wastes is degraded during composting (Zhang et al., 2019), and the large amounts of refractory cellulose and lignin continue to decompose slowly during the cultivation, which increases the aeration porosity. Second, with the growth and development of the seedlings, the root elongation squeezes the substrate and thus decreases the total porosity and water-holding porosity. Third, the growth of seedlings consumes large amounts of organic matters, nutrients, and minerals, which decreases the contents of organic matters, nitrogen, P_2_O_5_, K_2_O and total nutrients. The development of the seedlings also accelerates the decomposition of the refractory substances, resulting in the increases in EC value. The seedlings of group C show the highest survival rate, greatest seedling height and largest ground diameter (Table 6), and the aeration porosity of the substrate is increased by up to 104.05%. The total porosities of substrates A, B, C, D and E decrease by 21.35%, 13.31%, 7.95%, 5.38% and 20.11%, respectively, after the cultivation. Both substrates C and D contain palm fiber residues with high lignin contents (Table 1). The lignin is difficult to be degraded, which explains the smaller changes in the total porosity of the substrate. Therefore, it can be concluded that the porosity change of substate after seedling cultivation is related to the cellulose and lignin contents of the compost feedstock. Substrates A, B, C and D all contain Camellia oleifera shells with high hemicellulose contents (Table 1). The compost in substrate A was abstained in the absence of livestock and poultry manure and the microbial abundance (Jinping et al. 2021) and temperature of the compost were lower than those of the compost in substrate B (Figure 1). Therefore, the hemicellulose degradation rate of substrate A is lower than that of substrate B (Zhang et al. 2021). The degradation of hemicellulose in substrate A is continued during the cultivation of seedlings, resulting in the largest decrease in total porosity. The cellulose content of pine chips in substrate E is high (Table 1), which is more difficult to decompose, but easier than lignin. Therefore, small amounts of hemicellulose, cellulose and lignin continue to decompose, leading to the significant decrease in the total porosity of substrate E.

**Table 5.** Changes in the nutrients of compound substrates after seedling cultivation

| group |  | A | B | C | D | E |
| --- | --- | --- | --- | --- | --- | --- |
| organic | before | 54.33±2.72 | 50.90±2.53 | 43.00±2.13 | 37.00±1.82 | 61.50±3.06 |
| matters % | after | 9.45±0.46 | 19.14±0.94 | 14.43±0.73 | 9.93±0.48 | 15.76±0.77 |
|  | decrease | 82.61±5.85 | 62.40±6.82 | 66.44±6.65 | 73.16±6.22 | 74.37±3.51 |
| N% | before | 0.86±0.041 | 0.93±0.042 | 2.29±0.09 | 1.38±0.065 | 0.85±0.039 |
|  | after | 0.49±0.022 | 0.51±0.023 | 0.68±0.032 | 0.65±0.031 | 0.24±0.011 |
|  | decrease | 43.02±1.98 | 45.16±2.04 | 70.31±3.32 | 52.90±2.39 | 71.76±3.41 |
| P<sub>2</sub>O<sub>5</sub>% | before | 0.36±0.017 | 0.56±0.026 | 0.90±0.042 | 0.59±0.027 | 0.62±0.029 |
|  | after | 0.095±0.0045 | 0.16±0.0079 | 0.58±0.025 | 0.22±0.0099 | 0.069±0.0032 |
|  | decrease | 73.61±3.33 | 71.43±3.21 | 35.56±1.56 | 62.71±2.71 | 88.87±4.03 |
| K<sub>2</sub>O% | before | 1.47±0.074 | 1.68±0.082 | 2.13±0.097 | 1.43±0.068 | 0.75±0.035 |
|  | after | 0.59±0.027 | 0.72±0.034 | 0.63±0.031 | 0.61±0.029 | 0.43±0.021 |
|  | decrease | 59.86±2.86 | 57.14±2.74 | 70.42±3.43 | 57.34±2.73 | 42.67±1.87 |
| Total | before | 2.69±0.12 | 3.17±0.14 | 5.32±0.25 | 3.40±0.18 | 2.22±0.12 |
| nutrient % | after | 1.18±0.06 | 1.40±0.07 | 1.89±0.09 | 1.47±0.07 | 0.75±0.03 |
|  | decrease | 56.13±6.32 | 55.84±6.62 | 64.47±6.39 | 56.76±7.35 | 66.22±7.21 |

**Table 6.** Morphologies of seedlings raised on different substrates

| Treatment |  | A | B | C | D | E |
| --- | --- | --- | --- | --- | --- | --- |
| survival rate % |  | 99 | 99.5 | 100 | 97.5 | 99.5 |
| height/cm | average | 20.0±0.95 | 27.2±1.33 | 31.4±1.55 | 29.6±1.47 | 22.2±1.08 |
|  | Maximum | 44.0±2.10 | 51.0±2.46 | 63.0±3.05 | 47.0±2.23 | 38.0±1.85 |
| Ground | average | 2.8±0.13 | 3.7±0.17 | 4.1±0.22 | 3.9±0.18 | 3.3±0.15 |
| diameter/mm | Maximum | 3.9±0.18 | 4.7±0.22 | 5.5±0.26 | 5.1±0.23 | 4.4±0.21 |
| total root length/cm |  | 2169.79±107.96 | 4377.77±215.32 | 2161.92±107.21 | 2864.55±142.53 | 3900.97±192.67 |
| Root surface area/cm<sup>2</sup> |  | 371.30±18.23 | 768.81±38.23 | 397.98±19.56 | 596.43±29.62 | 683.89±34.01 |
| Average root diameter/mm |  | 0.56±0.026 | 0.56±0.027 | 0.58±0.027 | 0.66±0.032 | 0.56±0.025 |
| Root volume cm<sup>3</sup> |  | 5.10±0.24 | 10.86±0.52 | 5.90±0.30 | 10.01±0.51 | 9.58±0.45 |

### 3.4 Relationship between physical and chemical properties of compound substrate and seedling morphology

Survival rate is an important indicator for evaluating the successful cultivation of camellia oleifera seedlings. The survival rates of the seedlings of groups A, B, C, D and E are 99%, 99.5%, 100%, 97.5%, and 99.5%, respectively (Table 6), which are all above 97% and significantly higher than that (42.50-67.50%) obtained using the compost of rice hulls, bark, and sawdust (rice hulls 20%-26%, bark 28%-44%, sawdust 36%-52%) as the substrate (Lu et al., 2017).

Seedling height and ground diameter are important indicators for measuring the quality of seedlings (Wang et al., 2021). As shown in Table 6, the average heights and maximum heights of the seedlings cultivated on the 5 substrates for 6 months are in the order of C> D>B>E>A and C>B>D>A>E, respectively. Both of their average ground diameters and the maximum ground diameters are in the order of C>D>B>E>A. The average ground diameter (4.1 mm) and average seedling height (31.4 cm) of the seedlings in group C reaches the first-grade and second-grade criteria, respectively, defined in the standard GB/T 26907-2011 for two years old seedling (≥0.35 cm and ≥ 25 cm). Among them, 26.67% of the seedling heights reach the first-grade criteria (≥40 cm), and 45% of them reach the second-grade criteria. The average ground diameters of both groups B (3.7 mm) and D (3.9 mm) reach the first-grade criteria. Upon average seedling heights (27.2 cm and 29.6 cm), 6.7% of group B and 15% of group D reach the first-grade criteria and the rest all reach the second-grade criteria. The average ground diameter (3.3 mm) of the seedlings of E group meets the second-grade criteria ( ≥ 0.30 cm), and only 28.3% of the average seedling heights meet the second-grade criteria. Neither of the average ground diameter (2.8 mm) nor the average seedling height (20.0 cm) of the seedlings in group A conforms to the second-grade criteria, but 41.67% of the seedlings meets the second-grade standard for ground diameter and 15 % of the seedlings meet the second-grade standard for height. It is worth noting that the average seedling height of the seedlings in group C (31.4 cm) is 30.5 cm higher than those of two-year-old camellia oleifera seedlings cultivated on coconut bran and yellow soil (coconut bran 75% +yellow soil 25%) mixed with 3 kg/m^3^ of slow-release compound fertilizer from KOCH (USA) (Wang et al., 2018). The average seedling height (31.4 cm, 29.6 cm) and average ground diameter (4.1 mm, 3.9 mm) of the seedlings in groups C and D are better than those (28.70 cm, 3.93 mm) of the seedlings raised on the substrate formulated with peat and the compost of rice hulls (peat 50% + (compost of rice hulls+ extruded perlite + vermiculite + boiler slag + imported slow-release fertilizer) 50%) for 18 months (Cui et al., 2017). The seedling heights (27.2 cm, 31.4 cm and 29.6 cm) and ground diameters (3.7 mm, 4.1 mm and 3.9 mm) of the seedlings in groups B, C and D are better than those (25.98 cm, 3.43 cm) of the seedlings raised on the substrate containing composted bagasse, cassava skin and charcoal ash (2:1:1) for 18 months (Dai et al., 2016).

The compound substrate C contains the highest amounts of N, P_2_O_5_, K_2_O, and total nutrients before seedling cultivation, and the decreases in the contents of N, K_2_O, and total nutrient and the increase in aeration porosity are the most significant after the seedling cultivation (Table 5 and 6). The seedling survival rate, height, and ground diameter of this group are also the highest. These results suggest that seedling height and ground diameter are mainly related to the total nutrient content in seedling substrate, consistent with the results reported by Wu et al. and Zeng et al. (Zeng et al., 2021; Wu et al., 2018). In addition, the N and K_2_O contents of substrate C decrease by 70.31% and 70.42% respectively, which are the largest among those of the 5 substrates, while the decrease in its P2O5 content is the smallest, suggesting that nitrogen and potassium can promote the growth and development of camellia oleifera seedlings (Chen, 2009; Wang et al., 2010; Peng, 2019).

The organic matter contents of substrates A-E decreased by 82.61%, 62.40%, 66.44%, 73.16% and 74.37%, respectively, after the seedling cultivation due to the consumption by the seedlings and losses during watering. Before the seedling cultivation, the organic matter contents and total nutrient contents in the substrates are in the order of E>A>B>C>D, and C>D>B>A>E, respectively. The substrate with a high organic matter content generally has a low total nutrient content. The raw materials of the composts in substrates C and D are the same, but their formulas and ratios with other inorganic matrices are different. The portion of the compost in substrate C is 20% higher than that in substrate D. Therefore, both contents of organic matter and total nutrient of substrate C are higher than those of substrate D. The seedling heights and ground diameters of groups B, C, and D are significantly higher than those of the other two groups, but the decreases in their organic matter contents are smaller, indicating that there are large amounts organic matters in the composts of Camellia oleifera shell and livestock/poultry manures that can be absorbed and utilized by the seedlings. The maximum decrease in the total nutrients of substrate C is only 3.43%, and the average seedling height and ground diameter of the seedlings cultivated on it are the largest. Even though the organic matter content of substrate A is decreased by 82.61%, it shows no significant advantages for the growth of Camellia oleifera seedlings. It may be explained with the low degradation rate of lignocellulose during the composting process that results in low contents of organic matter usable for the seedling development. All these results suggest that the height and ground diameter of Camellia oleifera seedlings raised on the substrates are closely related to the composition of composting raw materials, composting formula, composting process and rate, and substrate formula. Therefore, stable composting raw materials, formula, composting process, and compound substrate formula are the key factors to improve the quality and efficiency of the cultivation of Camellia oleifera seedling.

The physical and chemical properties of substrate also affect the morphological characteristics of Camellia oleifera seedling root system. The root morphology can be characterized with the root growth indicators, such as root length, root surface area, root volume, and root diameter (Pang et al., 2018; Li et al., 2020). In general, the longer and thicker root systems with larger surface areas and volumes suggest better development and growth conditions. As shown in Table 6, the root morphologies of the seedlings raised on different substrates are significantly different. The seedlings of group B showed the longest root system of 4377.77 cm, the largest root surface area of 768.81 cm^2^, and the largest volume of 10.86 cm^3^, and both root surface area and volume of group A are the smallest. The root surface areas and root volumes of the seedlings raised on the five substrates are in the orders of B>E>D>C>A and B>D>E>C>A, respectively. The average root diameter of group D is 0.66 cm, significantly larger than those of other groups (0.56 cm for groups A, B and E, and 0.58 cm for group C). Despite the larger root length, surface area and volume of group B, its average root diameter is similar to those of other groups. Overall, the root development of group B is the best, possibly because the porosity and nutrient content of the substrate is more suitable for the root development of Camellia oleifera seedling. Yet the total root lengths, surface areas, and volumes of all five groups are better than those of the Camellia oleifera seedlings cultivated on traditional substrates, such as yellow soil, mushroom residue and peat substrate (40% yellow soil + 15% pine forest topsoil + 20% mushroom residue + 20% peat + 5% manure) for two years which show the average total root length of 280.32 cm, the surface area of 106.72 cm^2^, and the average root volume of 8.71 cm^3^ (She et al., 2020).

### 3.5 Correlation analysis of seedling Morphology and substrate physical and chemical properties

Seedling height, ground diameter and root activity are generally considered to be the most important indicators for evaluating the quality of seedlings (Lu et al., 2002; Ge et al., 2006). We selected seedling height, ground diameter and root morphology to evaluate the growth of Camellia oleifera seedling and analyzed the correlation between the physical and chemical properties of substrate and the seedling growth. As can be seen from Table 7, the moisture content of substrate is negatively correlated with seeding ground diameter (r=-0.972, P<0.01) and seedling height (r=-0.997, P<0.01). Seedling height and ground diameter are positively correlated (r=0.982, P<0.01), and the root surface area and root length are positively correlated (r=0.971, P<0.01). The total porosity of substrate before seedling cultivation is negatively correlated with seedling height (r=-0.891, P<0.05) and ground diameter (r=-0.944, P<0.05). The changes in the organic matter and total nutrient contents of substrate after seedling cultivation reflect the amounts of substrate nutrients absorbed by the seedlings and the nutrient losses of the substrate itself. The change in organic matter content is negatively correlated with ground diameter (r=-0.905, P<0.05) and seedling height (r=-0.957, P<0.05), and is positively correlated with moisture content (r=0.966, P<0.01), aeration porosity before seedling cultivation (r=0.952, P<0.05) and organic matter before seedling cultivation (r=0.906, P<0.05). The change in total nutrient content is positively correlated with the total nutrient content before seedling cultivation (r=0.983, P<0.01). The total nutrient content after seedling cultivation is negatively correlated with the aeration porosity before seedling cultivation (r=0.937, P<0.05) and is positively correlated with the total nutrient content before seedling cultivation (r=0.935, P<0.05). No correlation is found between the change in total nutrient content and seedling ground diameter (r=0.732) and seedling height (r=0.773). There are no significant correlations between organic matter and total nutrient contents before and after seedling cultivation and seedling ground diameter and height (r1=-0.723, r2=-0.823, r3=0.304, r4=184; r5=0.754, r6=0.820, r7=0.723, and r8=0.830). These results suggest that organic matters and nutrients that can be absorbed and utilized by Camellia oleifera seedling play a key role in the seedling growth and development.

**Table 7.**
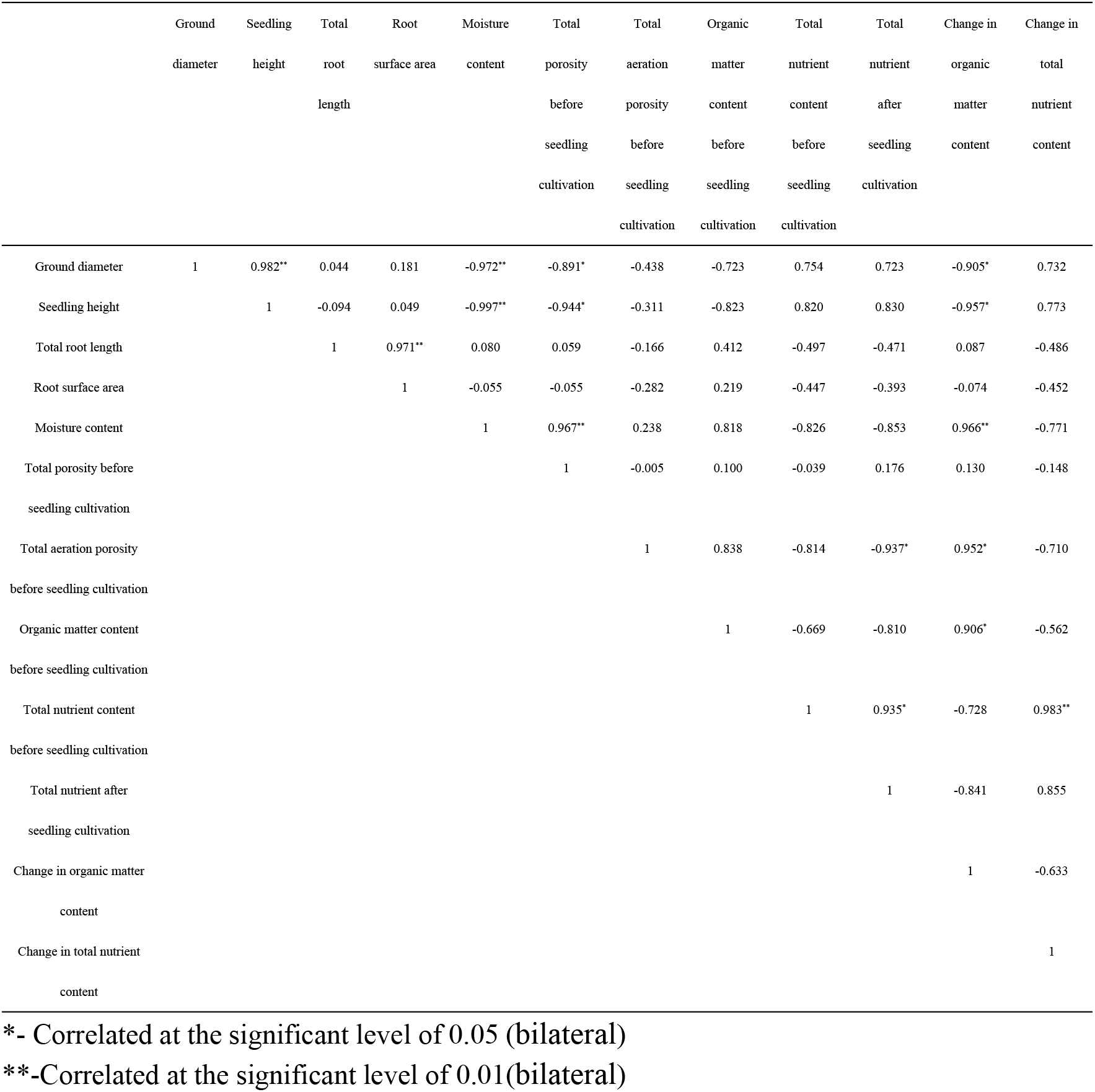
Pearson correlation analysis of physical and chemical properties of substrate and seedling morphology

## 4 Conclusion

Agricultural and forestry residues were composted using different formulas and the composting efficiencies were evaluated. It is found that the introduction of livestock and poultry manures and C/N ratios in the range of 25-35 can significantly increase the heating rate, prolong the thermophilic phase, and produce the composts with high total nutrient contents and GI.

The annual camellia oleifera cutting seedlings were cultivated after the composting products of agricultural and forestry residues were 100% substituted for peat and formulated with inorganic substrates including vermiculite and perlite. The physical and chemical properties of the substrates before and after seedling cultivation and the seedling development were analyzed. The results show that, first, the physical and chemical properties of composting raw materials, composting formula and degree of composting directly affect the physical and chemical properties of the obtained compost and corresponding compound substrate, as well as the growth of seedling. Second, the physical and chemical properties of compound substrate conform to the standards defined in GB/T 33891-2017 for seedling substrate, except that their pHs (7.03-7.43) do not meet the pH requirement (5.0-6.5) of the regulation LYT 2314-2014. However, the Camellia oleifera seedlings developed better on these substrates than on traditional nursery substrates. Therefore, further study is needed to evaluate the effects of substrate pH on the growth of Camellia oleifera seedling. Third, the survival rates of the seedlings on all five substrates are ≥97.5% after 185 days cultivation covering the sever hot summer and cold winter. In particular, the seedlings cultivated on substrate C that contains the highest amount of total nutrient reaches exhibited 100% survival rate, and the largest seedling height and ground diameter. In addition, the seedling heights and 26.67% of ground diameters reach the first grade criteria for two years old seedling. Forth, the changes in the physical and chemical properties of substrate after seedling cultivation directly affect the growth and development of Camellia oleifera seedling. The initial total porosity is negatively correlated with seedling height at the significant level, and the change in organic matter content is negatively correlated with ground diameter and seedling height, and positively correlated with the initial aeration porosity and initial organic matter content at significant levels. It is then concluded that the amounts of organic matter and total nutrients in the substrate that can be absorbed and utilized by the seedling play a key role in the seedling development.

## Acknowledgements

The authors are grateful for the financial support from the National Key R&D Program of China (Grant No. 2019YFD1001602) and the Provincial Department of Science and Technology of Zhejiang, China(Grant NO.2017C02022).

## References

[1] Dai X, Sun W, Fan Q, Luo P, He J. Physicochemical property of mixed substrates with agricultural and forestry wastes and comprehensive evaluation of their effect on growth of Camellia oleifera seedlings. Journal of Plant Resources and Environment.2016; 25(1):54–61.https://doi.org/10.3969/j.issn.1674-7895.2016.01.07

[2] Guo W, Qi W, Wang Z, Wang C. Analysis on the research status and development of agriculture and forestry waste resource utilization under the condition of blending and briquetting. Journal of China Agricultural University. 2021; 26(1):143–150.https://doi.org/10.11841/j.issn.1007-4333.2021.01.15

[3] Zhao Y, Liu J, Zhang B. Effects of Different Substrate Ratio of Agricultural and Forestry Waste and Slow-release Fertilizer on the Growth of Ornamental Pepper. Horticulture & Seed. 2021; 41 (09):6–9. https://doi.org/10.16530/j.cnki.cn21-1574/s.2021.09.003

[4] Lu H, Chen W, Su Y, Xie P. Research on fertilizer effect of forest waste compost as seedling substrate. Rural Economy and Technology.2021; 32(10): 17–20.

[5] Nie L. Research on the present situation and development strategy of Agricultural waste resource recovery in China. Agricultural Economic Research. 2017; 07: 59–60.

[6] Wu Q, Chen J, Gan F, Li C, Jiang H, Zheng H. Application of Well-composted Sawdust and Litterfall Substrate to Camellia oleifera Seedling. Guangxi Forestry Science.2017; 46(4): 431–435. https://doi.org/10.19692/j.cnki.gfs.2017.04.021

[7] Luo J, Tan X, Peng Y. A Study on the Composting Technology for Camellia Shell Medium. Acta Agriculturae Universitatis Jiangxiensis. 2011; 33 (4):0712–0718.https://doi.org/10.13836/j.jjau.2011127

[8] Zhuang R. China camellia. Beijing: China Forestry Publishing House, 2008.

[9] Wang M, Zhang Y, Tang X. Report of Light Media Nursery Test For Camellia. Forestry and Environmental Science.2018; 34(6): 56–60.

[10] Zhang X, Ding X, Zhang Y, Liu Y, Cai J, Li Y. Effects of nutrient deficiencies on root system in Camellia gauchowensis seedlings. Nonwood Forest Research.2014; 32(4): 170–174. https://doi.org/10.14067/j.cnki.1003-8981.2014.04.033

[11] Wang Q, Peng G. Influence of substrate and time on cuttage steckling of Camellia oleifera. Forest By-Product and Speciallity in China. 2021; 04: 001–003.https://doi.org/10.13268/j.cnki.fbsic.2021.04.001

[12] Li Y, Ma J, Wang D, Wei Y, Ye H. Effects of Mediums with Different Physicochemical Properties on Cutting Rooting of Camellia osmantha. Chinese Journal of Tropical Agriculture. 2020; 40(3):25–30. https://doi.org/10.12008/j.issn.1009-2196.2020.03.005

[13] Lv B, Jiang M, Ye F, Zheng X, Yu S. Research on matrix formula of oil tea container seedling. South China Agriculture. 2017; 11 (5):24–26.https://doi.org/10.19415/j.cnki.1673-890x.2017.05.014

[14] Yang L, Chen W, Chen X, Chen J, Weng C, Zheng D. Effects of different light substrates and containers on rooting of camellia oil cuttings and evaluation of matrix bagging methods. Modern gardening. 2018; 10:43–45. https://doi.org/10.14051/j.cnki.xdyy.2018.19.022

[15] Liu C, Li J, Shu Q. Growth Comparison of Camellia oleifera Seedlings Indoor and Outdoor with Different Substrates. Guangxi Forestry Science. 2019; 48(3):332–335.https://doi.org/10.19692/j.cnki.gfs.2019.03.010

[16] She Y, Zhang Y, Cao K, Li X, Zhang L, Wang Y. Effects of different culture medium formulas on roots biomass and morphology of Camellia oleifera container seedling. Non-wood Forest Research. 2020; 38 (4): 11–16. https://doi.org/10.14067/j.cnki.1003-8981.2020.04.002

[17] Hua M, Wang J. Experimental instruction of soil physics. Beijing: Beijing Agricultural University Press. 1993;61–73.

[18] GB/T 33891-2017, Organic substrates for greening. Beijing: Standards Press of China, 2017.(in Chinese)

[19] Shuang L, Ni S, Du J, Teng Z, Shi C. Determination of Organic Carbon in Geochemical Soil Sample by Potassium Dichromate Oxidation-Heating Method. Anhui Chemical Industry. 2016; 42(4): 110–112.

[20] Bremner JM Nitrogen-total. In: Sparks, D.L. (Ed.), Methods of Soil Analysis. Part 3 – Chemical Methods. SSSA Inc., ASA Inc., Madison, WI, USA, pp. 1996;1085–1122.

[21] Tiquia SM, Tam NFY and Hodgkiss IJ Effects of turning frequency on composting of spent pig-manure sawdust litter. Bioresource Technology 1997; 62(1-2): 37–42.

[22] Wang Y, Li M, Qiu H, Zhang W, Zhang C, Li Y. Changes of microbial quantity and nutrient content in different composting of livestock manure. Journal of Gansu Agricultural University. 2017; 52(3), 37–45. https://doi.org/10.13432/j.cnki.jgsau.2017.03.007

[23] Qiao, X, Shen, G, Wang, Z, Guo, C, Liu, Y, Lei, Z, Zhang, Z. Co-composting of livestock manure with rice straw: Characterization and establishment of maturity evaluation system. Waste management, 2014; 34(2):530–535. https://doi.org/10.1016/j.wasman.2013.10.007

[24] Guardia, A.D, Mallard, P, Teglia, C, Marin, A, Pape, C.L, Launay, M, Benoist, J.C, Petiot, C. Comparison of five organic wastes regarding their behaviour during composting: part 1, biodegradability, stabilization kinetics and temperature rise. Waste Management, 2010; 30(3), 402–414. https://doi.org/10.1016/j.wasman.2009.10.019

[25] Ma R, Li D, Qi C, Li G, Wang G, Liu Y, et al. Effects of C/N ratio on compost maturity and odor emission of chicken manure. Transactions of the Chinese Society of Agricultural Engineering. 2020; 36 (24):194–202. https://doi.org/10.11975/j.issn.1002-6819.2020.24.023

[26] Wang Q, Liu G, Zhang R, Li Y, Liu A, Wang W. Effects of Physical and Chemical Properties of the Substrate on the Growth of Container Seedlings of Ficus religiosa. Journal of Northwest Forestry University. 2021; 36 (5):88–93. https://doi.org/10.3969/j.issn.1001-7461.2021.05.1

[27] Wu J. Comparison of physical and chemical properties of several solid cultivation substrates. Journal ofJilin Agricultural Sciences. 2006; 31 (4):17–20.https://doi.org/10.16423/j.cnki.1003-8701.2006.04.006

[28] LYT 2314-2014. Technical specification for seed raising of oil tea containers. Beijing: State Forestry Administration, 2014. (in Chinese)

[29] Wu J, Li X, An S. Effects of Different Substrate Ratios on the Growth of Sour Pummelo Seedlings. Chinese Journal of Tropical Crops. 2018; 39 (3):443–447. https://doi.org/10.3969/j.issn.1000-2561.2018.03.006

[30] Zeng L, Li J, Wu F, Huang S, Li M, Cai Y. Effects of different substrate formulations on substrate nutrient content and seedling growth of Macadamia nut. Northern Horticultur. 2021; 24: 51–56. https://doi.org/10.11937/bfyy.2021135

[31] Chen H. Studies on factors concerned in nutrient media aeration improving growth as well as yield in cucumber. Shenyang,Liaoning, Shenyang Agricultural University, 2009.

[32] Wang Y, Shao X, Huang X, Wang K. Advances in studies on nitrogen uptake in plant roots. Pratacultural Science. 2010; 27 (7): 105–111.

[33] Peng Z. Absorption, transportation and regulation of nitrogen element in plants. Journal of Hebei Agricultural University. 2019; 42 (2):1–5. https://doi.org/10.13320/j.cnki.jauh.2019.0024

[34] Ma S. Effects of Application Amount and Fertilization Modes of N,P and K on Growth,Yield and Quality of Eggplants. Shandong, Shandong agricultural University, 2018.

[35] Li C. Effects of Nitrogen,Phosphorus and Potassium Fertilization on the Growth and Physiological Characteristics of Saraca Dives Seedlings. Guangxi, Guangxi University, 2020.

[36] Yin J. Effects of Nitrogen Rate and Ratio of Nitrogen,Phosphorus,and Potassium on Dry Matter Accumulation and Grain Filling of Castor Bean (Ricinus Communis L.)

[37] Lu M, Jiang F, Song X. Assessing indices of container seedling quality. Chinese Journal of Applied Ecology. 2002; 13 (6):763–765. https://doi.org/10.13287/j.1001-9332.2002.0181

[38] Ge Y, Zhu J, Jiang B, Yuan W, Shen A. Study on evaluation index of container seedling quality. Journal of Zhejiang Forestry Science and Technology. 2006; 26 (1): 10–12, 22.

[39] Wang X, Lin X, Li Y, Ruan Y, Xing X. Physical and chemical properties of several kinds of agriculture and forestry waste composite matrix and their effect on container seedling of Phoebe chekiangensis. Journal of Zhejiang A & F University. 2013; 30 (5):674 – 680.

[40] Li J, Zhao X, Wei S, Zheng J, Hu Y. Study on the Physico-Chemical Properties of soil-Less Cultural Substrates of Pollution-Free Vegetable. Journal of Southwest Agricultural University. 2000; 22 (2):112–115. https://doi.org/10.13718/j.cnki.xdzk.2000.02.006

[41] Pang S, Ma Y, Zhang P, Lao Q, Yang B, Liu S. Effects of Substrate Ratio and Applying Dosages of Slow-release Fertilizer on the Growth of Container Parashrea chinensis Seedlings. Journal of Northwest Forestry University. 2018; 33 (6):66–71.https://doi.org/10.3969/j.issn.1001-7461.2018.06.11

[42] Peng S, Chen Y, Yang X, Wang X, Ma L, Wang R, et al. Study on the Classification of Seedling Quality of Oil-tea Camellia (Camellia oleifera). Chinese Agricultural Science Bulletin. 2012; 28 (25): 101–105.

